## Supporting Information for "Computational characterization of the xanthan gum glycosyltransferase GumK"

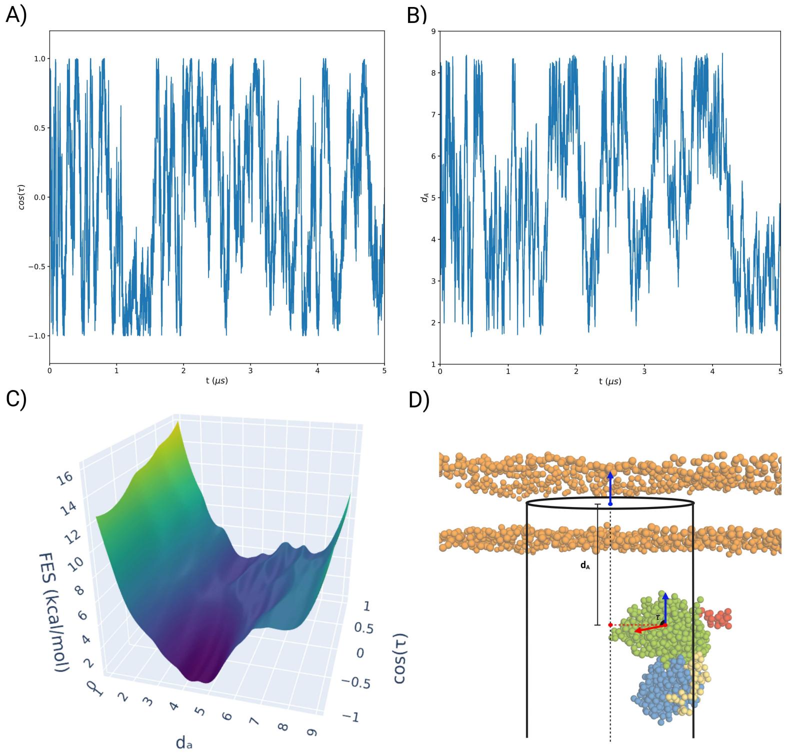


**Fig S1.** Quality assessment of the cylindrical-restrained OPES simulation using the protein–protein rescaled-interaction MARTINI force field. A) Time evolution of the tilt angle of the acceptor domain relative to the membrane. Multiple transitions are sampled during the simulation. B) Time evolution of the distance between the acceptor domain and the membrane. Several transitions are observed, and when the distance falls below 4 nm, the protein is considered bound. C) Three-dimensional reweighted free-energy surface (FES) showing two main minima. D) Schematic representation of the cylindrical restraint. The black cylinder illustrates a $z$-aligned positional restraint applied to a reference atom in the acceptor domain. The membrane distance $d_A$ is the projection of this atom onto the cylinder axis, while the tilt angle $\tau$ is measured relative to the membrane normal ($z$-axis) using a second acceptor-domain atom to define the internal vector.

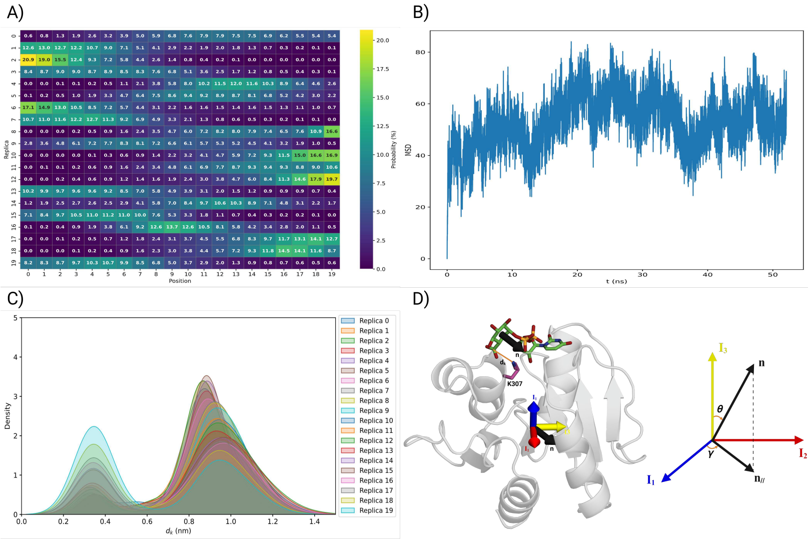


**Fig S2.** Quality analysis of the Hamiltonian replica exchange simulation used to sample glucuronate-UDP bound to the GumK donor domain. A) Probability matrix of the 19 replicas involved in the simulation. The replicas generally diffuse along the ladder, although some remain trapped in specific positions. B) Mean square displacement of the replicas in replica space. This indicates that while the replicas diffuse well along the ladder over certain periods, the system does not reach full stationarity, suggesting that the simulation is not fully converged. C) Distribution of the distance between the sugar C6 atom and the $N^{\zeta}$ atom of Lys307. The results show that the replicas sample two main states: one where the carboxylate group interacts with Lys307 (below 0.6 nm) and another where the phosphates interact with the carboxylate. D) Definition of the polar space used to describe the orientation of the sugar ring relative to the protein. The normal vector of the sugar ring is computed and translated to the origin of the inertia axes of the $\alpha$-carbons in the initial donor-domain conformation. The entire trajectory is first aligned to this structure to ensure that the inertia axes remain consistently aligned across all frames. The polar angles of the translated normal vector are then used to describe the orientation of the sugar ring. The distance between C6 and Lys307 is also shown.

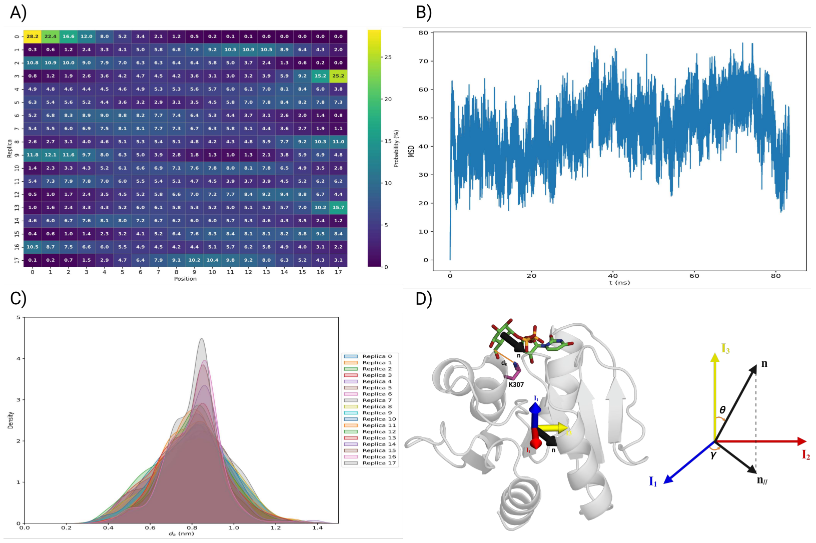


**Fig S3.** Quality analysis of the Hamiltonian replica exchange simulation for glucose-UDP bound to the GumK donor domain. A) The probability matrix shows good mixing of the replicas, with only a few remaining trapped in specific positions along the replica ladder. B) Mean square displacement of the replicas in replica space shows stable behavior, with replicas diffusing efficiently. However, a decrease in the trend at the end of the simulation suggests that full convergence may not have been reached. C) Distance distribution between the sugar C6 atom and the $N^{\zeta}$ atom of Lys307. The distribution is multimodal but consistently centered around 0.8 nm across all 17 replicas. This distance is compatible with the phosphate groups interacting with Lys307. The same definitions of the collective variables (CVs) as in Figure S2 are used in this case.

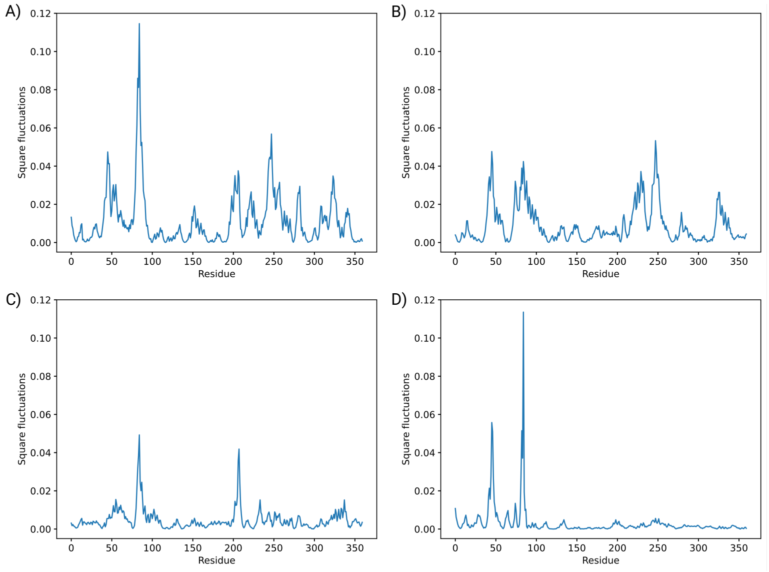


**Fig S4.** Square fluctuations of the first four normal modes derived from the normal-mode analysis of the GumK crystal structure. A) The first normal mode (NM1) involves coordinated movements across different regions of both domains, primarily corresponding to a twisting motion. B) The second normal mode (NM2) exhibits smaller squared displacements compared to NM1 but still engages both domains, mainly reflecting a bending motion. C) The third normal mode (NM3) shows squared displacements similar to those of NM2, with motions becoming more localized within each domain. This mode helps define the opening path described in the main text. D) The fourth normal mode (NM4) is highly localized within the acceptor domain, indicating that it primarily describes local backbone vibrations rather than large-scale conformational changes.

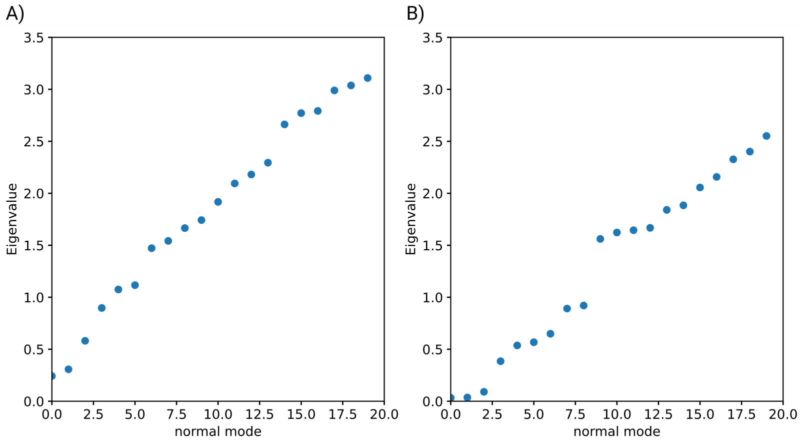


**Fig S5.** A) Eigenvalues of the first 20 normal modes computed from the normal-mode analysis of the GumK crystal structure (PDB: 2HY7). Several discontinuities are observed, but the first two modes are closely clustered, whereas the third mode begins to exhibit a higher frequency. B) Eigenvalues of the first 20 normal modes of the open conformations sampled by ClustENMD. The first three normal modes approach zero, suggesting nearly free relative motion of the two domains.

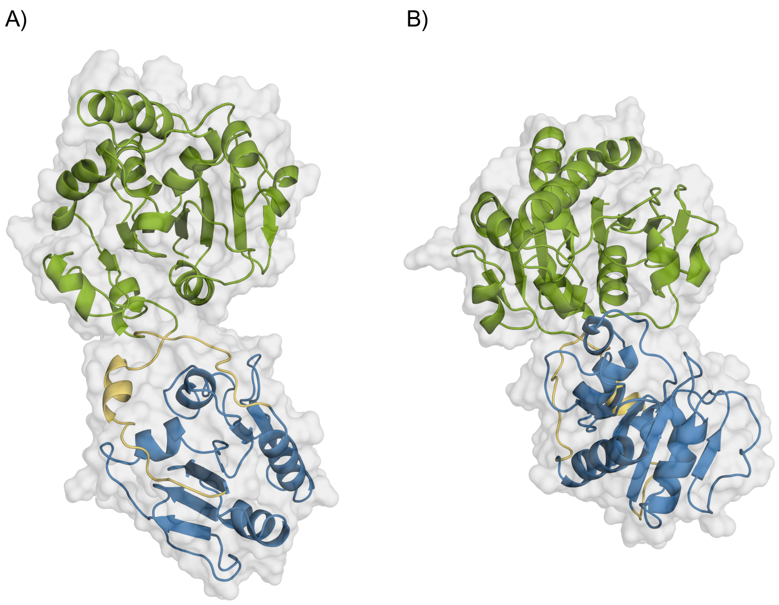


**Fig S6.** Sampled conformations from the ClustENMD simulation. A) An overtwisted conformation located at the periphery of the ensemble in the 3D normal-mode space. The donor domain is rotated by 180° around the main axis due to the high flexibility of the yellow loops. B) A metastable state positioned at positive values of normal mode 3. Helix C$\alpha$4 occupies the acceptor substrate-binding site, likely representing an inactive conformation that is disfavored when the acceptor substrate is bound.

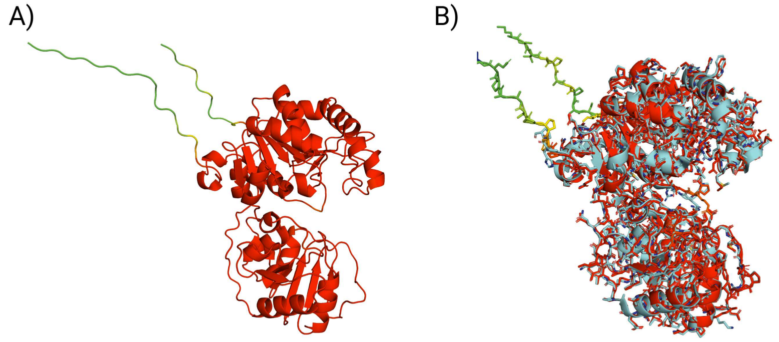


**Fig S7.** Analysis of the AlphaFold-predicted structure of GumK. (A) The structure is colored according to the pLDDT score, showing low confidence only in the terminal regions. (B) Superposition of the AlphaFold model with the crystal structure (PDB ID: 2HY7), showing that both the backbone and the side chains closely match the experimental structure, with RMSD values of 0.4 Å and 0.5 Å, respectively.

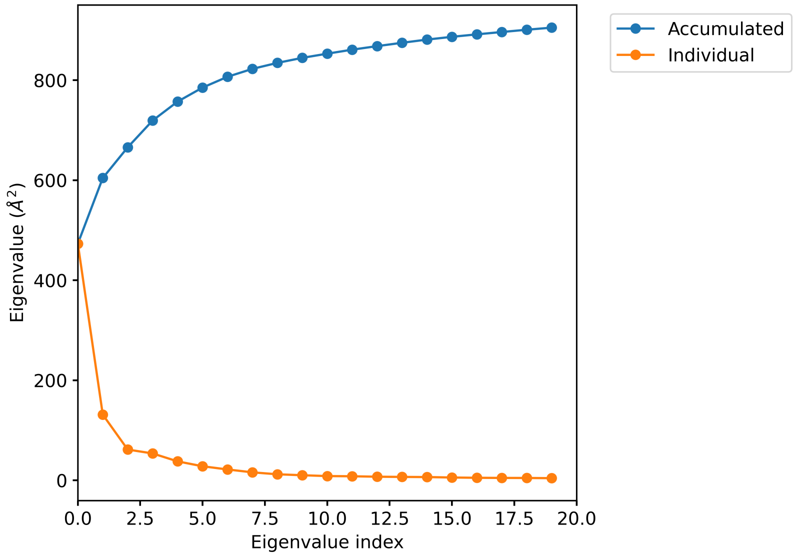


**Fig S8.** Variance of the first 20 principal components from the essential dynamics analysis of the unbiased MD simulation of GumK in the membrane-bound state. The first five principal components are sufficient to define the essential subspace, as the individual variance decreases sharply after the fifth component and approaches an asymptotic value. The cumulative variance shows that the first few principal components account for a significant percentage of the total variance.

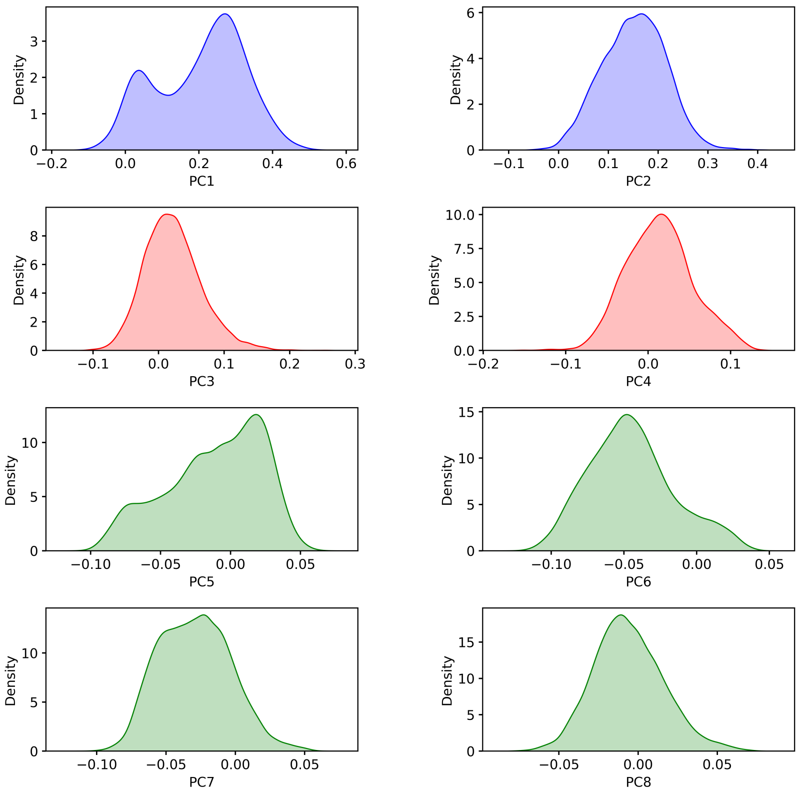


**Fig S9.** Probability distributions of the trajectory frames projected onto the first eight principal components (PCs). The first principal component exhibits a well-defined bimodal distribution, suggesting a possible transition pathway. In contrast, the last principal component displays a Gaussian distribution, indicating that this direction behaves harmonically or corresponds to a fast, fully sampled motion. Based on these distributions and the eigenvalue analysis, seven principal components are sufficient to define the essential subspace of GumK.

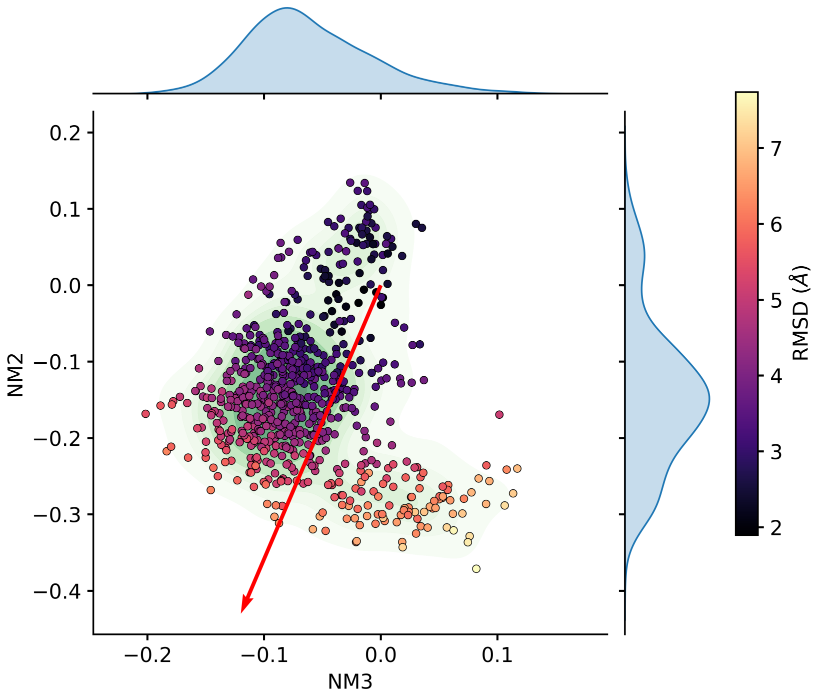


**Fig S10.** ClustENMD ensemble exploring a region near the GumK crystal structure using a target RMSD of 0.5 $\mathring{A}$. The distribution in the space defined by normal modes 2 and 3 shows that the protein follows a path close to that defined by the first principal component from the essential dynamics analysis of the unbiased MD simulation. The red arrow represents the projection of the first principal component onto the normal mode space.

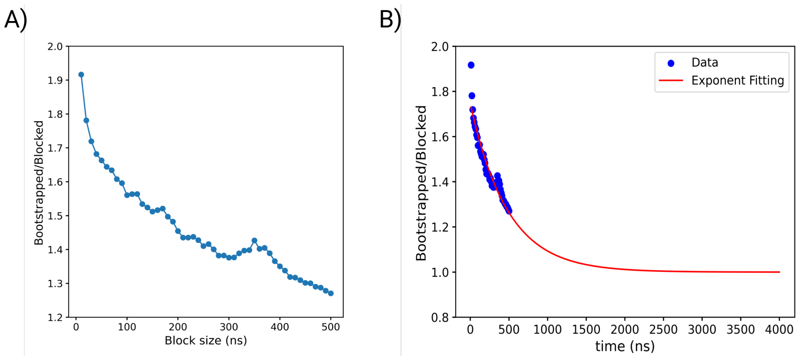


**Fig S11.** A) Bootstrap-normalized block covariance overlap of a 1 µs simulation of GumK in the membrane-bound state. The curve is expected to decay exponentially toward 1 as the essential subspace becomes fully sampled. B) Exponential fit of the decay, indicating that at least 3 µs of simulation are required to achieve satisfactory convergence.

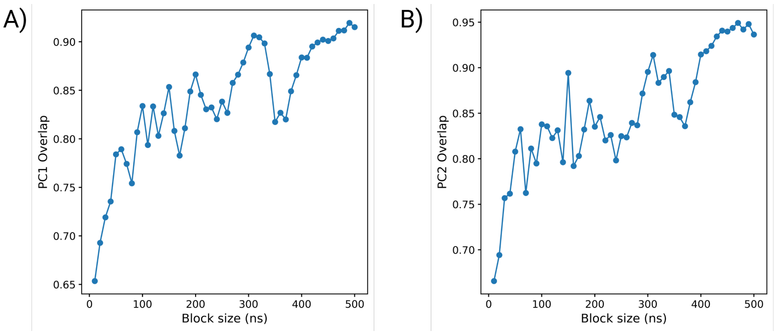


**Fig S12.** Overlap analysis of the first and second principal components (PCs) with their final counterparts at different time points. After 400 ns, when partial opening is sampled, the directions of the two PCs show good overlap with the final ones.

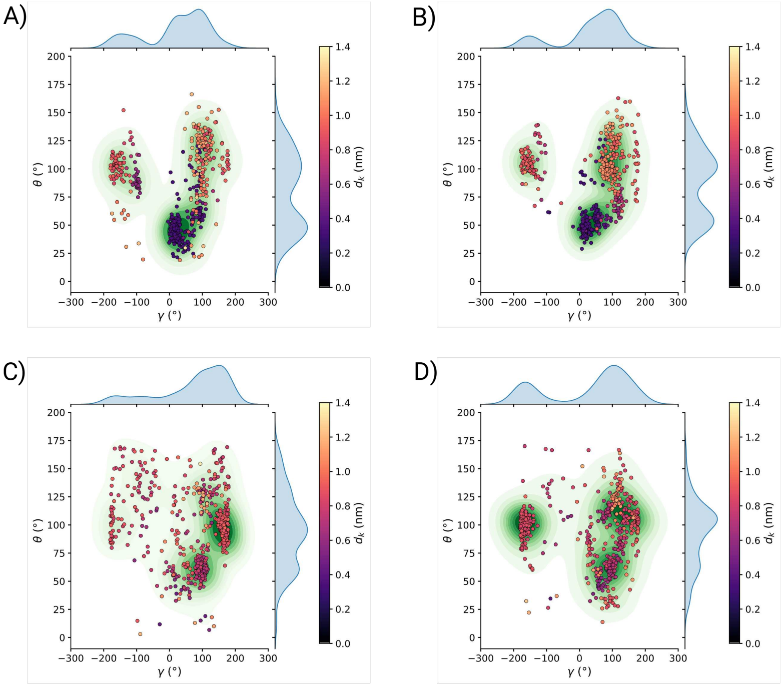


**Fig S13.** Distribution of hydrophobic interaction conformations in the GumK donor domain for the four clusters sampled during the Hamiltonian replica exchange simulation of glucose-UDP (Fig. 6 in the main text). In all cases, the hydrophobic interactions are more dynamic than those observed for UDP-glucuronate (Fig. S15).

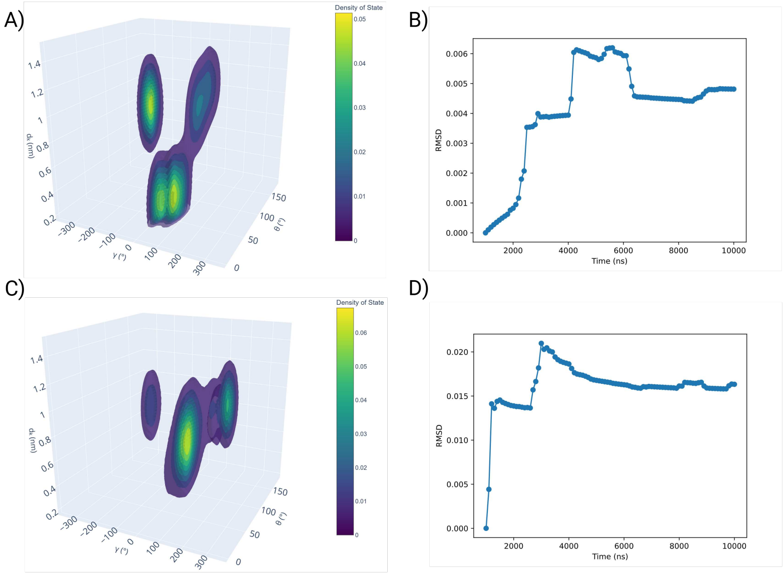


**Fig S14.** A) Probability density of glucuronate-UDP as a function of three collective variables (CVs): the distance between the C6 atom of the sugar and the $N^{\zeta}$ atom of Lys307, defined as $d_k$, and the two angles $\gamma$ and $\theta$ describing the orientation of the sugar ring in a polar coordinate system. B) RMSD of the probability density with respect to the initial state, calculated after 1 ns of simulation. The density of states is complex, and stable populations are observed at the end of the simulation. C) Probability density of glucose-UDP. D) RMSD of the density of states calculated as in B). In this case, convergence is even clearer. The density functions were computed by including all replicas from the Hamiltonian replica exchange simulation, with weights determined using the WHAM algorithm.

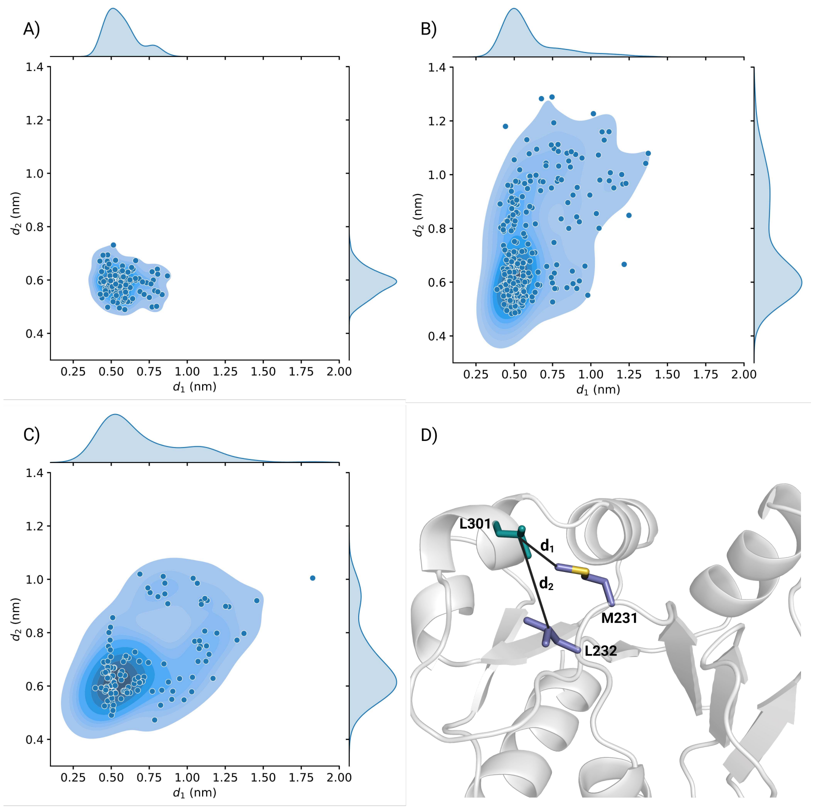


**Fig S15.** Distribution of hydrophobic interaction conformations in the GumK donor domain for the three clusters sampled during the Hamiltonian replica exchange simulation of glucuronate-UDP. (A–C) Panels A, B, and C correspond to clusters A, B, and C, respectively, as defined in Figure 6 in the main text. Cluster A exhibits a well-localized distribution, with the residues consistently remaining in contact. In contrast, clusters B and C display more dynamic interactions, showing greater variability in residue contacts. (D) Definition of the two distances used to characterize these distributions.

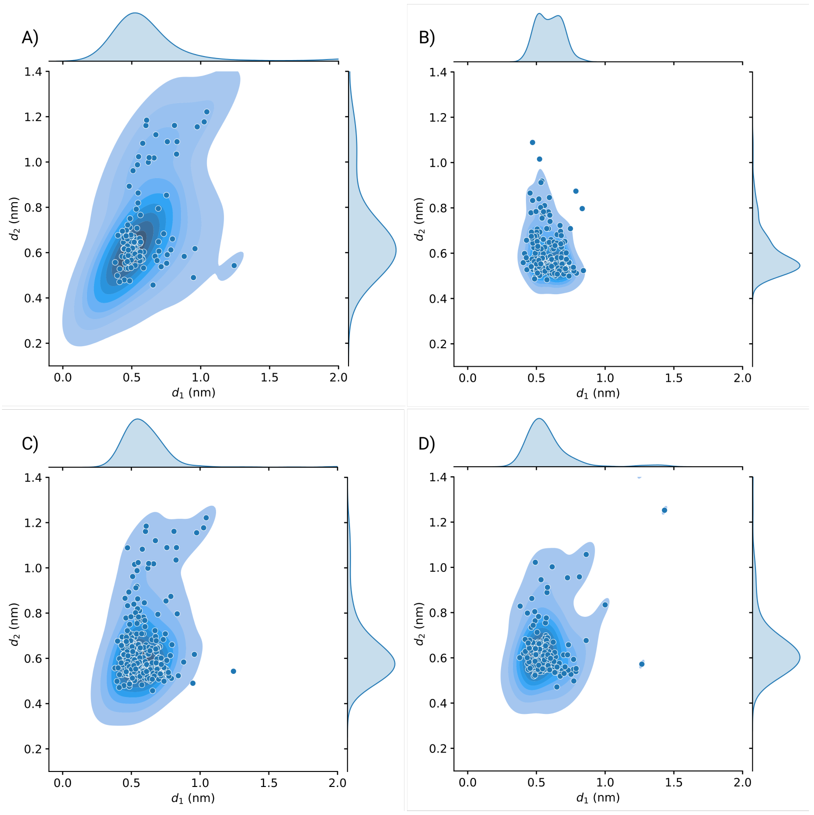


**Fig S16.** Distribution of hydrophobic interaction conformations in the GumK donor domain for the four clusters sampled during the Hamiltonian replica exchange simulation of glucose-UDP (Fig. 6 in the main text). In all cases, the hydrophobic interactions are more dynamic than those observed for glucuronate-UDP (Fig. S15).

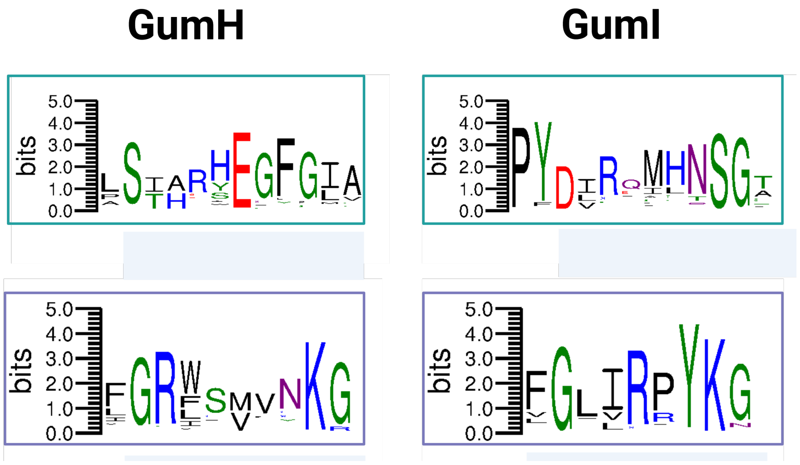


**Fig S17.** Structural alignment of the regions of GumI and GumH that are expected to interact with the sugar portion of the donor domain, i.e., Mannose-GDP. The color scheme follows the same convention as in GumK (Fig. 3 in the main text). The distribution of positive charges is reversed compared to GumK, with two positive charges present in the purple region, which is neutral in GumK. Additionally, the green region is shorter than in GumK, where it includes a loop with a highly variable sequence between residues Ala293 and Pro298.

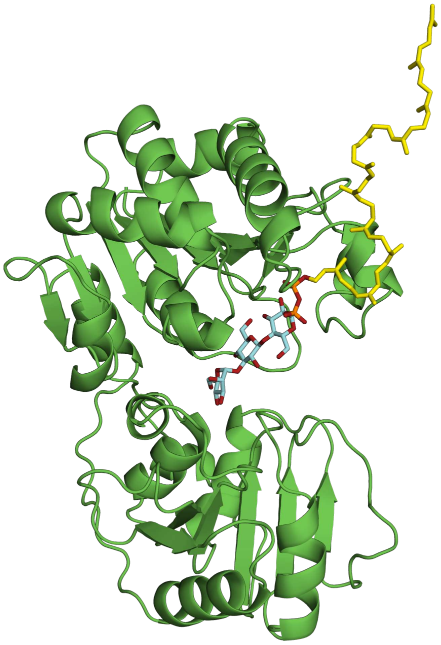

**Fig S18.** Starting conformation of the GumK–acceptor complex. The substrate was manually positioned near the binding site based on insights obtained from the unbiased MD simulations of the free protein in solution and from previous test simulations.

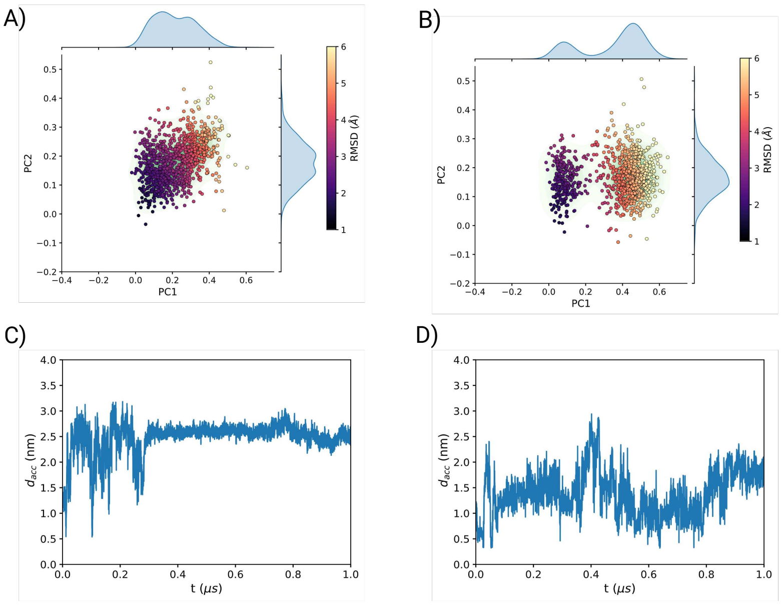


**Fig S19.** (A, B) Projections of replicas 1 and 2 onto the first two principal components. In the presence of the acceptor substrate, GumK undergoes conformational changes, although not to the same extent as in the main simulation (see main text). (C, D) Distance between the reactive oxygen atom of the acceptor substrate and the catalytic residue Asp157. In these replicas, the acceptor substrate does not establish stable interactions as observed in the main simulation, which explains why the conformational space of the protein appears more restricted.

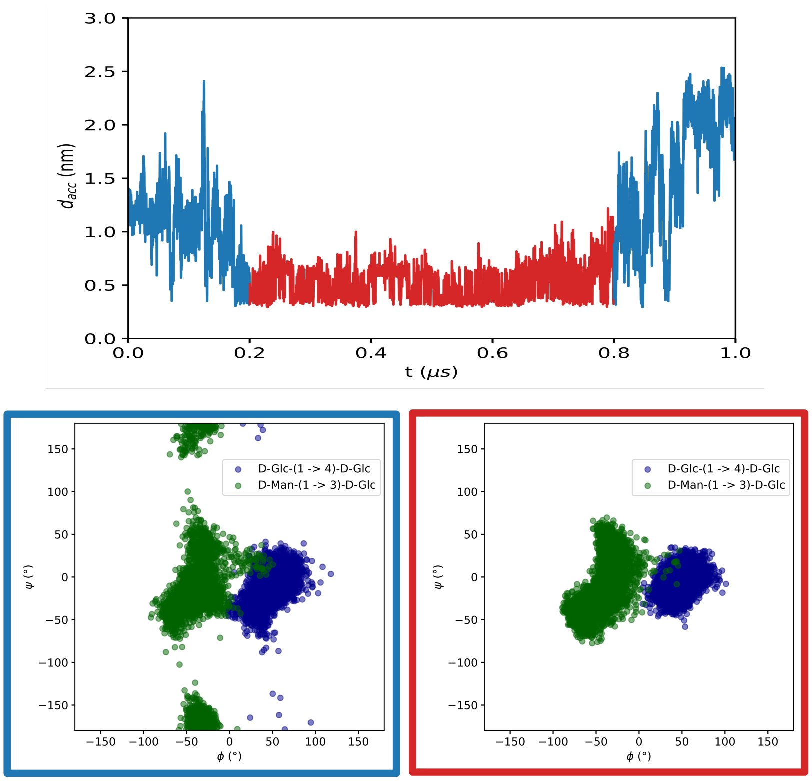


**Fig S20.** Distance distribution of the reactive hydroxyl group of the acceptor substrate relative to the catalytic residue Asp157. Below, the Ramachandran plots show the distributions of the two glycosidic bonds in the acceptor substrate of GumK (Fig. 1B) sampled during the simulation. The red color represents the bound state. The distributions indicate that when the sugar is bound, it is constrained to a specific conformation corresponding to the fully extended state. The blue dots represent the glycosidic bond between the first two glucose units in the acceptor substrate. This bond is more localized than the glycosidic bond between the second glucose and mannose, suggesting that in the bound state, the first unit is more restricted, while the second remains more flexible as it is exposed to the cleft between the two domains.

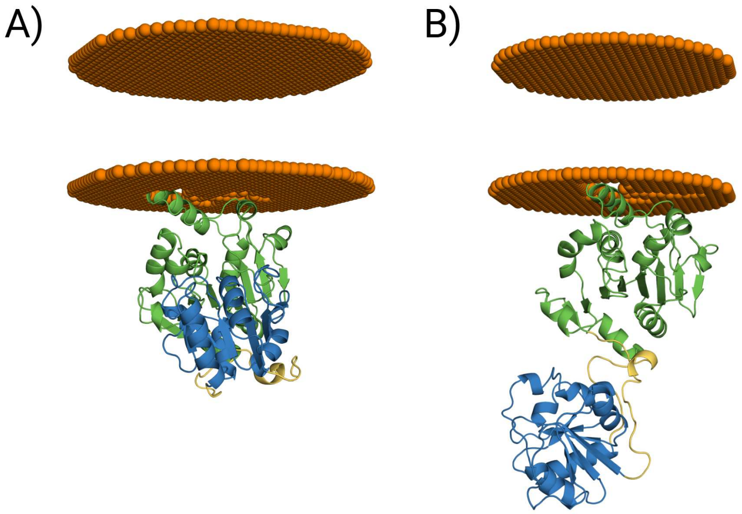


**Fig S21.** A) OPM prediction of the GumK–membrane complex with the protein in an close conformation. Helix N$\alpha$4 is inserted into the membrane through its two tryptophan residues. B) OPM prediction of the open conformation of GumK interacting with the membrane. The conformation was taken from ClustENMD and shows a binding mode similar to panel (A), except for minor differences due to the distinct conformation of the clamp.

**Table S1.** This table reports the functional classification of the residues identified in the simulation.

| **Region** | **Residue** | **Function** |
|---|---|---|
| Donor domain | Lys307, Tyr292, Ser305, Ser230, Arg29 | Sugar diphosphate interaction |
| | Leu301, Met231, Leu232, Met306 | Steric constrain |
| Acceptor domain | Leu56, Val89, Phe92, Lys60, Arg96 | Tail interaction |
| | Glu131, Ser22, Ser132, His23, Arg52 | Sugar interaction |
| Interdomain | Arg29, Glu272 | Opening |
| | Tyr328, His201, Ser160, Glu192, Asp157, Thr161, Asp303, Ser186, Met189, Ser304 | Twisting |
